# Bioisosterism reveals new structure-odor relationships

**DOI:** 10.1101/567701

**Authors:** Narmin Tahirova, Erwan Poivet, Lu Xu, Zita Peterlin, Dong-Jing Zou, Stuart Firestein

## Abstract

The question of structure-odor relationship (SOR) has inspired numerous studies into chemical odorants via their perceptual similarities. Much of this data comes from psychophysical studies on humans, precluding the possibility of direct measurements of receptor or receptor neuron activation. Remarkably, we find that many of the perceptual classifications used by human subjects translate well enough to mouse that we can apply cellular methods to better understand the molecular mechanism, that leads to odor discrimination and perception. Using a well studied and well recognizable odor percept of bitter almond, we have tested an odorant panel of aldehydes and ketones that were reported to share such perceptual qualities. These odorants include aromatic and aliphatic, as well as cyclic and allylic features. When parsing these odorants using chemical descriptors, we have a clear separation of molecules possessing these various features. However, here we show that OSN responses better recapitulate the physiological percept. Using these odorants, we also provide a proof-of-concept for non-classical bioisosterism at work in the olfactory system.

## INTRODUCTION

The mammalian olfactory system discriminates between a vast array of at least millions of organic chemicals we term odors. The majority of these odors are organic molecules with molecular weights below 350 Daltons and displaying many of the physical chemical characteristics common to small organic molecules [1]. One problem in understanding the chemistry of olfaction is that many of the same chemical features describe molecules that have no odor, to us and frequently to other animals tested. Over decades various attempts to classify odors based on chemistry and psychophysics (e.g., perceptual labels such as fruity, woody, fishy, etc.) have failed in one way or another, and none have provided a rational method for explaining why an odor has a particular quality – or not.

We have been pursuing a method of classifying odors that include biological as well as chemical and psychological parameters. This method is borrowed from the pharmaceutical industry and is generally called medicinal chemistry. It is based on using chemistry and biology to discover what are termed bioisosteres – combining the idea of isosteric chemical structures with similarity in biological function [2]. In this case however the chemical sterism is not necessarily based on the same parameters as an organic chemist, interested in synthetic reaction centers for example, would use. Thus two odors that had similar perceptual qualities but different molecular structures might still be considered bioisosteres – the key being that different molecular structures than those of interest to an organic chemist maybe the isosteres.

In previous work we have looked at the roles of heteroatomic ring substitutions and esters which have a particularly malleable structure. In both cases we found that the biological responses could be grouped according to relatedness and these groupings differed from those that a chemical classification program produced [3, 4]. In this paper we further examine the role of bioisostereism by focusing on non-classical bioisosterism, which introduces seemingly structurally discrepant groups as biological substituents. One often cited example of this is “ring opening” within a larger chemical structure, where medicinal chemists attempt to reduce the number of catalytic sites of a drug to improve its retention time by substituting a ring with a structurally rigid alkyl chain. Interestingly, odorants studied by Boelens in 1974 eliciting a bitter almond smell, an odor facet so well described that at one time it was been deemed the *model par excellence* for SOR studies in olfaction [1], contain non-classical bioisosteres with just such alterations. The best characterized molecule in almond extract is benzaldehyde, which contains a fully conjugated benzene ring attached to an aldehyde. Other iterations used by Boelens, however, are either devoid of aromaticity or even do away with any ring structure, while still maintaining some of the benzene’s perceptual qualities. The persistence of the percept is somewhat surprising given the significance assigned to electron conjugation and structural integrity of a ring in synthetic chemistry. With new tools to explore how benzene and its derivatives interact with the olfactory system, this study revisits the bitter almond odorants with a medicinal chemistry twist.

## RESULTS

### Dissociated OSN responses to aldehydes

Making use of medicinal chemistry strategies we utilized an odorant panel consisting of benzaldehyde (BEN) and several of its derivatives (Fig. 1), some of which are reported in the perfumery literature to elicit similar perceptual qualities (Fig. 2). The effect of aromaticity on odorant-OR interaction was tested using alkyl rings such as cyclohexene-1-carboxaldehyde (CYC6-1), cyclopentane carbaldehyde (CYC-5), and 3-cyclohexene-1-carbaldehyde (CYC6-3). The allylic odorants, such as tiglinaldehyde (TIG-4), 2-methyl 2-pentenal (TIG-5), and butyraldehyde (BUT), were used to probe the relationship between cyclic and non-cyclic odorant activation space. BUT was expected to be the outgroup due to the lack of a rigid skeleton in all of the other odorants. Acetophenone (ACE), a ketone which has only one additional carbon and shares perceptual descriptors with BEN, was also tested. For comparing these more similar odorants, an additional outgroup odorant, 2,3,5-trimethyl-3-thiazoline (TMT), was added. TMT is a component of fox urine and has been shown to elicit a fear response in rodents. Since odor responses projecting to amygdala are thought to be separate from other neutral or pleasant smells, there was good reason to believe that TMT would elicit a significantly different OSN response pattern versus the rest of the panel.

**Fig. 1.**
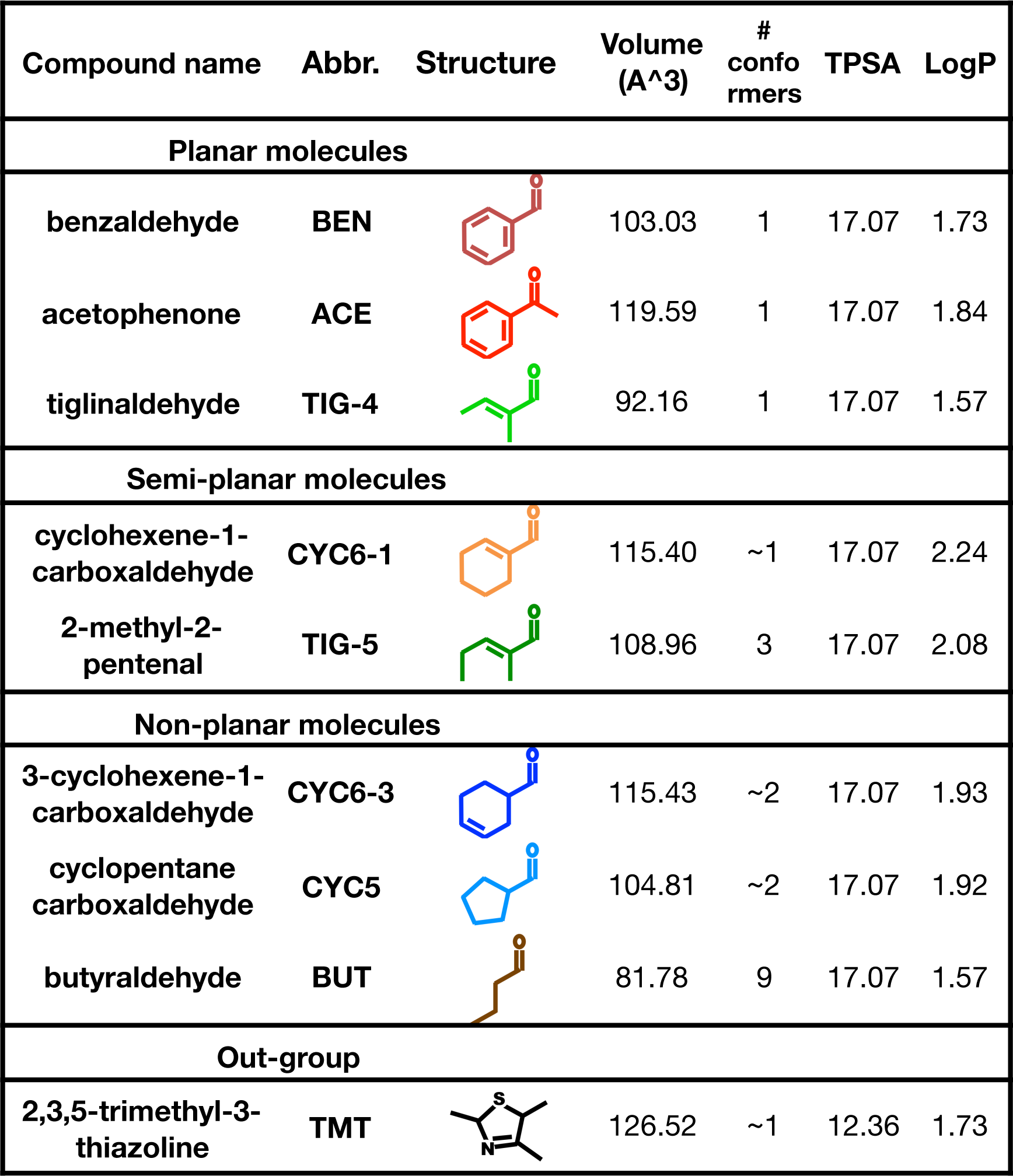
Benzaldehyde-derived odorant panel. Panel of aldehydes and ketones designed to study the relationship between the aliphatic, aromatic, and allylic odor spaces. Since TPSA has been shown to play a key role in discrimination of the odorants in the ketone and ester panels tested previously, we used a panel of odors with the same value as previously studied ACE. Planar molecules are rigid due to complete conjugation, and semi-planar molecules are only planar around the electronegative carbonyl group. All others assume a tetrahedral conformation at saturated single bonds. In alkanes, a simple rule of thumb is that each single bond multiplies the number of possible conformers by three. Cyclic alkanes are restricted to far fewer conformations, and aromatic rings can adopt just one. TMT, a component of the fox urine eliciting fear response in rodents is added as an out-group.

**Fig. 2.**
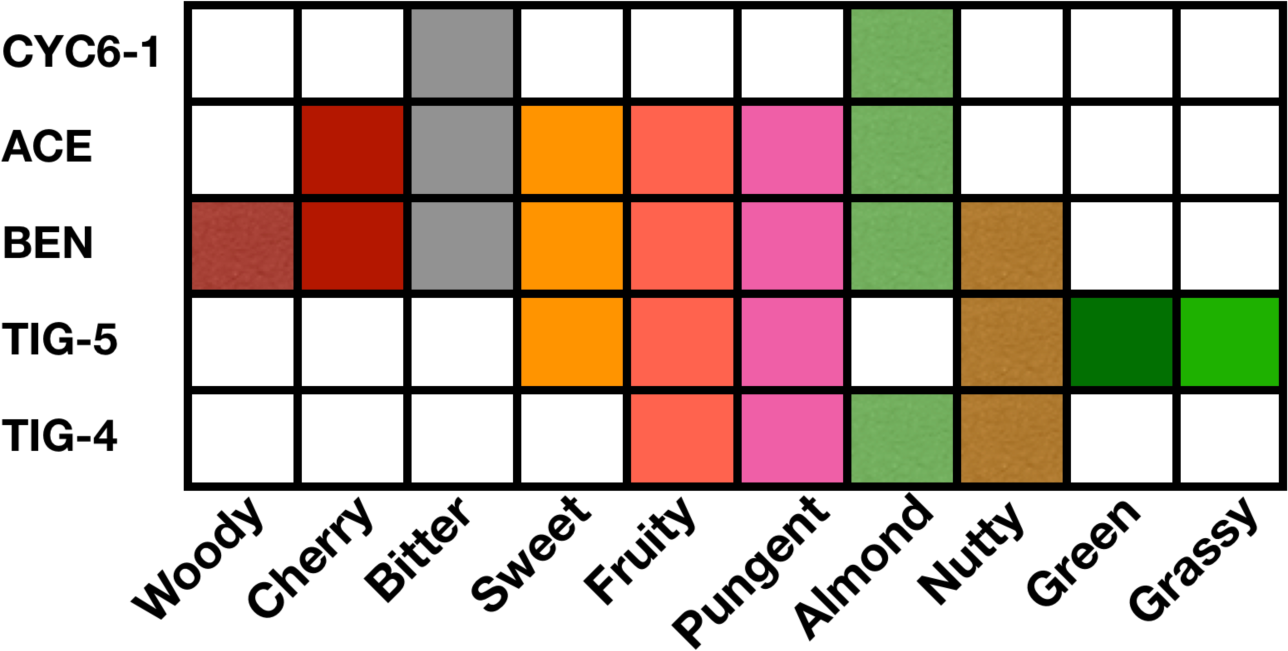
Perceptual overlap. Schematic of the psychophysical descriptors assigned to the panel odorants from www.goodscents.com. Some of these structures (CYC6-1, ACE, BEN, TIG5) have been used in SOR studies of the bitter almond accord. The absence of descriptors does not necessarily indicate the lack of those perceptual characteristics in the odorants.

Of 3027 live OSNs, 506 OSNs responded to at least one odorant in the panel. When the responses were scored in a binary fashion of “response/no response,” 68 response patterns emerged (Fig. S1). The number of patterns is typically taken to be the minimum number of OR types responding to the panel of odorants. It is possible to find differently tuned receptors within each response pattern type, where key features of the odorants likely play a bigger role in activation of specific receptors (Fig. 3), so we reiterate that this would represent the minimum number of receptors. Overall, 6.7% of live OSNs (204/3027) responded to BEN, 9.2% to CYC6-1, 3.1% to TIG4, 4.7% to TIG5, 6.2% to CYC5, 7.3% to CYC6-3, 4.8% to ACE, 4.7% to BUT, and 3.5% to TMT. The most highly activating odorant, CYC6-1, has the highest octanal to water solubility coefficient (LogP), while the odorant activating the least number of OSNs, TIG4, has the lowest LogP, with a loose correlation between this variable among the rest of the odorants (r=0.703).

**Fig. 3.**
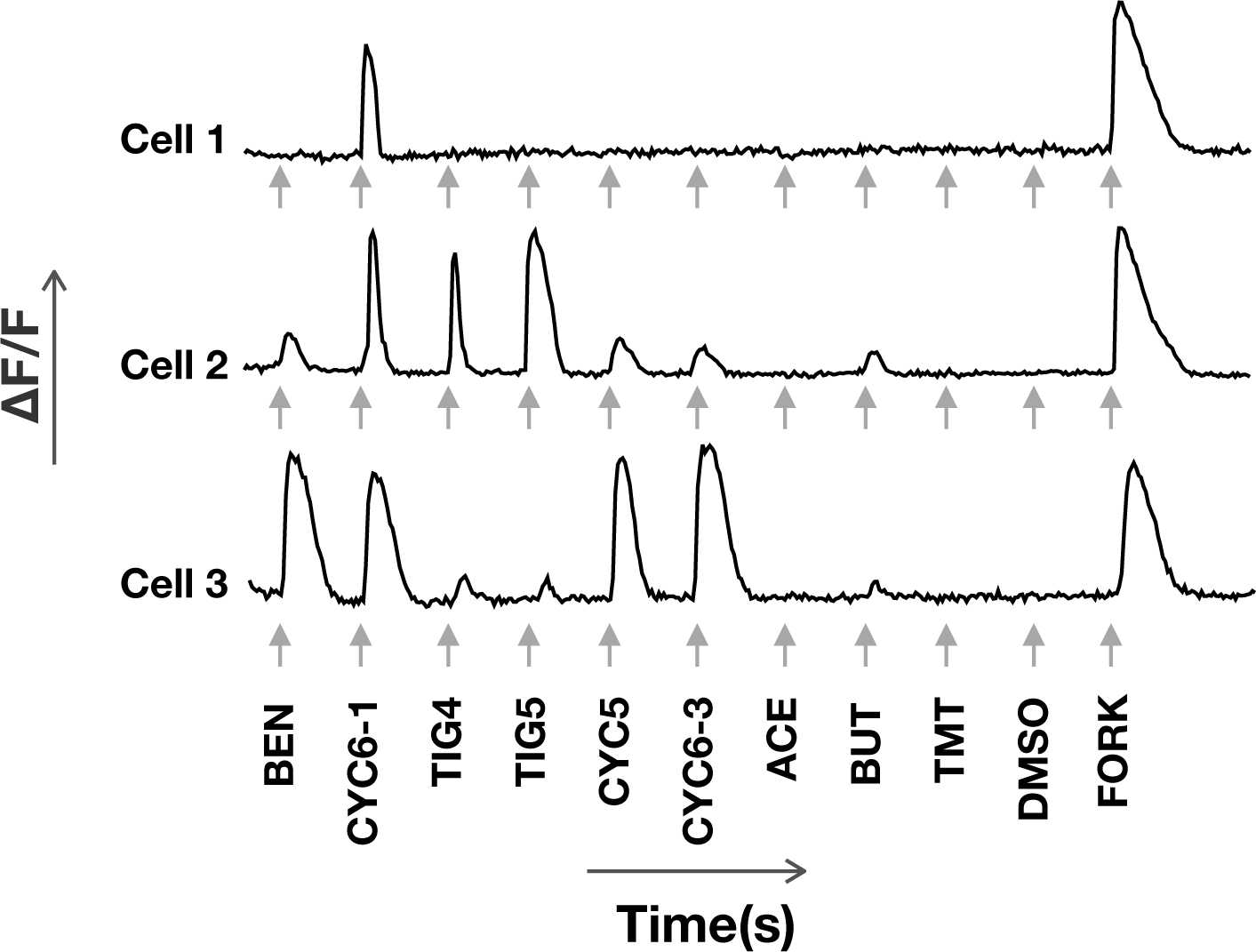
Calcium imaging traces. Sample traces of the OSN responses to 30uM odorant stimulations, where peak onset after stimulation (shown with arrows) indicates OR activation. The negative control contains equimolar concentration of dimethyl sulfoxide (DMSO), a polar aprotic solvent used to dilute all odorants. Forskoline (FORK) is an AC3 agonist used to probe cell viability is a positive control. Cell 1 is an example of a narrowly tuned cell. Cells 2 and 3 are activated by the same odorants, but show differential tuning to specific structural elements. While cell 2 responds more prominently to chemicals that contain a tiglic moiety, cell 3 responds strongly to cyclic aldehydes.

While 84% of the 204 OSNs responding to BEN also responded to CYC6-1, only 20% responded to TMT, the outgroup. 16 out of 506 responding OSNs (16/506) were activated by all of the odorants in the panel. 288/506 were only activated by the aldehydes, and 44 of these responded to all of the aldehydes. 199/506 were only activated by cyclic aldehydes and/or ACE, with a large activation overlap between the ketone and the aldehydes. The odorants with the highest exclusive activation were TMT, with 50 OSNs responding only to it, and BUT with 42 OSNs, both of which were originally included as “outgroup odorants”. Some patterns were observed in many OSNs, such as all cyclic aldehyde activation (seen in 23 OSNs), general aldehyde activation (seen in 20 OSNs), and all odorant activation (seen in 16 OSNs). 15 of the 69 patterns were only observed in one OSN each.

Dendrograms based on binary responses (Fig. 4) reveal a significant overlap between aliphatic, aromatic, and allylic odor spaces, as predicted by medicinal chemistry principles. The non-ring compounds, TIG4 and TIG5, that mimic the extracted portion of the aromatic rings clustered closely with BEN and ACE. The non-planar rings cluster farther away from the planar and semi-planar odorants, and a conformationally free BUT clustered the farthest out of the aldehydes. ACE clustered closely to the other aldehydes (Fig 4, Fig 5a), while TMT, which was chosen as the outgroup, did in fact cluster away from the rest of the panel, with barely any activation overlap. Interestingly according to the e-dragon chemical descriptors’classification, odorants were expected to segregate in aliphatic or cyclic molecules with cyclic molecules being segregated based their ring size (5 or 6 atoms) and finally on their aromaticity.

**Fig. 4.**
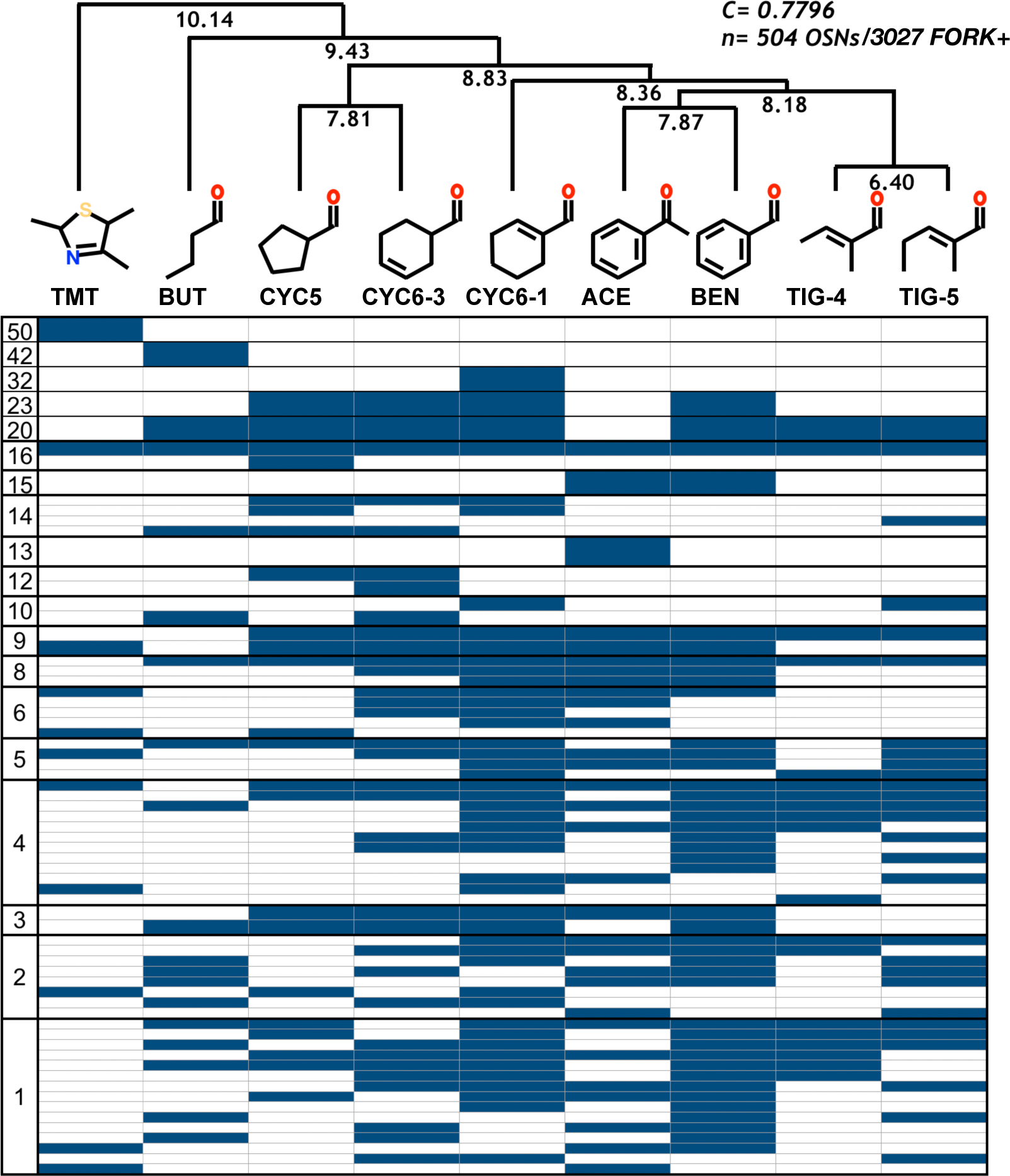
Responses of the OSNs to the panel. OSN responses from 504 OSNs are conservatively ranked in a binary fashion and represented in a clustered table, with each column corresponding to the odorants listed on the top. Each row in the diagram shows a distinct response pattern, and the number of OSNs responding in a certain pattern is tabulated on the left hand column. The dendrogram linking the Euclidean distances (numbers at the fork points of the dendrogram) of the odorants based on the hierarchical cluster analysis of the binary score is displayed at the top. *C* denotes the cophenetic correlation coefficient.

**Fig. 5.**
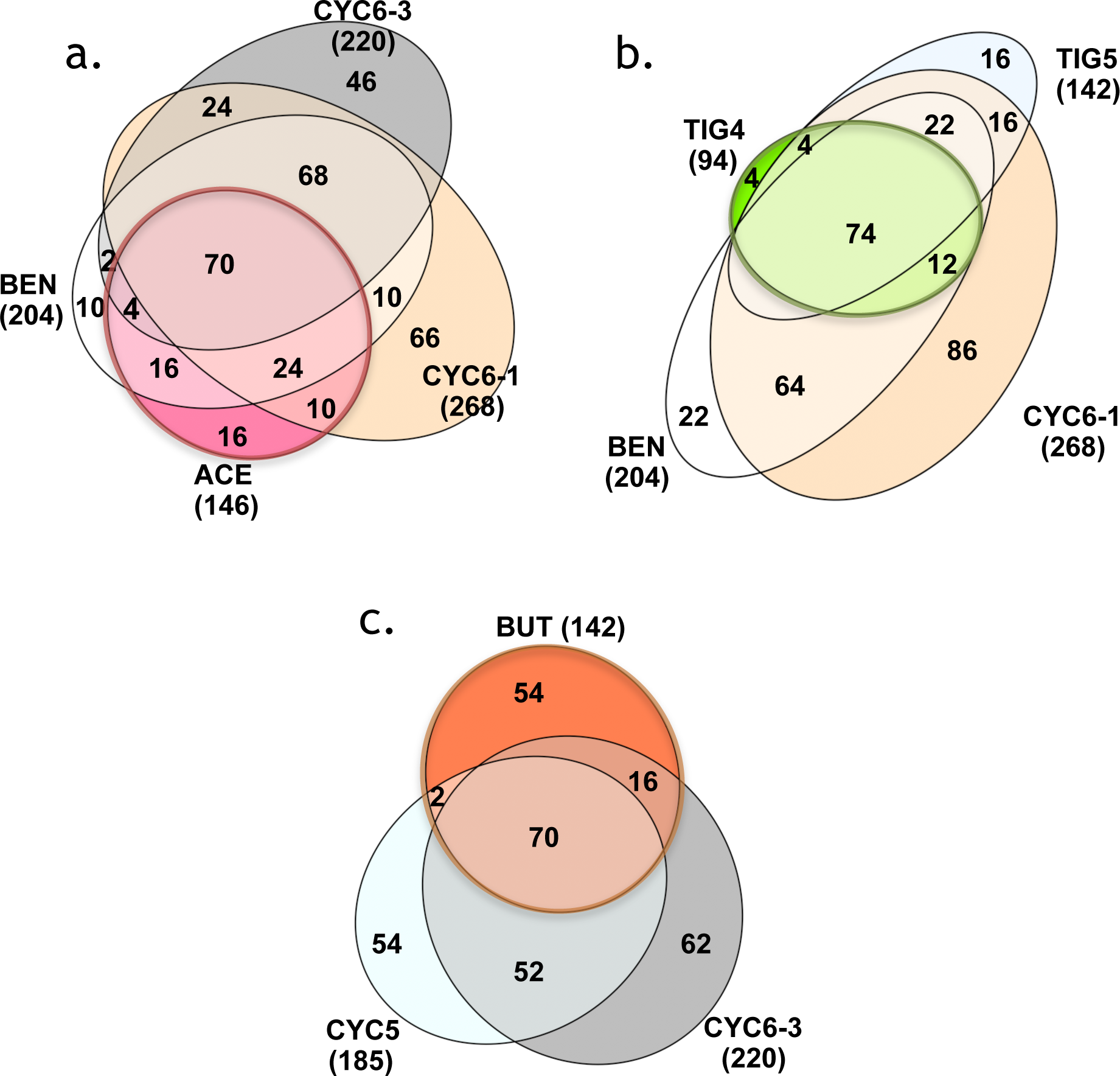
Venn diagrams representing odorant activation. **a.** A significant overlap can be seen between the ketone and aldehyde activation (e.g. ACE and BEN) and between aromatic and aliphatic activation (e.g. BEN and CYC6-1)” **b.** Odorants containing the tiglic moiety share a significant overlap. Most notably, conformationally restricted extraction of CYC6-1, TIG4, is almost entirely overlapping with the rest of the odorant, highlighting the role that conformational restriction plays in activation overlap. Similarly, TIG5 with an extra rotable bond leads to the recruitment of extraneous OSNs (16 cells in TIG5’s light blue compared to only 4 in TIG4’s bright green). **c.** BUT, which can shape into extracted portions of CYC6-3 (as well as CYC5), while overlapping similarly closely to the CYC6-1 and TIG4 pair, has a significant amount of extraneous activation (54 cells in orange), likely caused by conformational freedom afforded by two rotable single carbon bonds.

Examining the OSN activation overlap of the odorants sharing structural homology also revealed several interesting insights. Comparing the odorants containing a 6-membered ring (Fig 5a), the different structural elements that may affect binding can be seen. First, the so-called aromatic rings activate fewer OSNs than the aliphatic rings. It appeared that, their name notwithstanding, the fully conjugated aromatic rings did not in fact increase odor potency. There is also a significant overlap between the aliphatic and aromatic odorants, as well as non-aldehyde ACE greatly overlapping with the rest of the aldehydes. While many panels used in traditional literature on functional imaging of OSNs has categorized odorants based on their functional groups [3-8], our results suggest once again that functional groups should not be the primary distinguishing criteria for the odorants. CYC6-1 and CYC6-3, while sharing the chemical formula, TPSA, weight, and volume, show a surprisingly small amount of overlap, indicating a presence of a significant structural differentiator. The Jaccard similarity index is 55% among BEN and CYC6-1, 51% among BEN and CYC6-3, and 50% among CYC6-1 and CYC6-3, indicating that there is a roughly similar proportion of OSNs which are “broadly” tuned to some kind of an aldehyde with a six-membered ring. The Jaccard similarity index with ACE is 48.3% for BEN, 35% for CYC6-1, and 27% for CYC6-3.

The extent of aliphatic and aromatic activation space overlap can be appreciated when comparing compounds containing the tiglic moiety (Fig. 5b) as for example, with the odorants containing the cyclohexane structure (Fig. 5a) vs. the odorants containing the tiglic moiety (Fig. 5b). While smaller, less lipophilic odorants activate fewer OSNs, TIG4, a directly extracted fragment of CYC6-1 with a rigid structure, activated nearly all the OSNs which are part of the CYC6-1’s activation repertoire. Thus TIG4 is a strong bioisostere in a subset of OSNs responding to the larger compounds. Among the OSNs co-activated by both CYC6-1 and TIG4, the average calcium peak heights, adjusted to Forskolin (given a peak height of 1), were 0.79 and 0.38, indicating that TIG4 on average has a weaker potency.

Interestingly, in some of its conformations, BUT can act as an extracted fragment of CYC6-3 (Fig. 5c). 86 of 220 CYC6-3 responding OSNs are activated by BUT (39%), while 90 out of 268 CYC6-1 responding OSNs are activated by TIG4 (34%). Strikingly, while only 4 TIG4 responding OSNs don’t overlap with CYC6-1, 56 BUT responding OSNs are non-overlapping with CYC6-3. This additional activation by BUT is likely due to its conformational freedom, allowing it to access binding regions that a more rigid CYC6-3 structure may not reach. Similarly, with the TIG5 overlap with the cyclic compounds (Fig. 5b), there is an increased number of non-overlapping OSNs, which could be due to the conformationally free extra carbon.

### Odorant classification by chemistry or by OSN responses

We compared how the aldehydes in our panel are parsed using the traditional chemistry-centered approach and the OSN response patterns. Using the binary score of the OSN responses from 504 OSNs, we clustered the biological responses using hierarchical cluster analysis (Fig.4, Fig.S4a). Specifically among the aldehydes (Fig.S4a), the OSN response cluster highlights the structural elements found within the odorants. The structurally rigid odorants such as TIG4, TIG5, CYC6-1, and BEN form a cluster despite the discrepancy in size, presence of the ring, and electron conjugation properties. The semi-rigid CYC6-3 and CYC5 are later combined in this cluster, while conformationally free BUT forms an outgroup to the aldehydes.

While the OSN response dendrogram articulates the discrimination between rigidity of the molecule, clustering odorants based on all 1,666 chemical descriptors provided by the eDRAGON applet leads to an emphasis on separation between the higher molecular weight cyclic odorants and those with smaller chains (Fig. S4b). Specifically, the bond linking the carbon cycle to the electronegative functional group seems to be of importance in the OSN response cluster, while this is inconsequential in the full eDRAGON descriptor cluster as CYC6-1 and CYC6-3 are clustered together. Interestingly, when run with a full panel of odorants, the eDRAGON cluster also shows strong emphasis on full electron conjugation, as both ACE and BEN, which have a different polar functional group, cluster together.

In an attempt to reverse-engineer the OSN response patterns using the chemical descriptor database, we calculated and compared the distance matrix of every molecular descriptor provided by e-Dragon with our biological classification using their Spearman’s correlation factor (Table S1). In order to refine the descriptors that would best separate out the molecular responses in the aldehydes, we excluded TMT from this analysis. Our previous publication using the descriptor extraction has excluded the ketone from the analysis due to inability to recapitulate the OSN clustering pattern (Poivet et al 2018). In the present study, excluding ACE has significantly improved the rank correlation of the aldehydes. We therefore extracted the 10 highest ranking molecular descriptors (Table S1), and used them to reconstruct the odorant cluster (Fig. S4c). The highest ranking descriptor emphasizes the rigidity of the carbon attached to the aldehyde group, recapitulating the level of planarity in the panel’s aldehydes. Structural rigidity has also been cited in SOR study of sandalwood [9]. The most frequently seen type of descriptor, the edge adjacency index (ESpm), is used to define the topographic shape of the molecules.

### Behavioral response of mice to the odorants, and its relationship to the OSN activation

In order to better understand the implications of certain OSN response patterns to higher order processing of the stimuli, we used a simple pairwise odor comparison test in mice (Fig. S5). Briefly, mice were consecutively presented with a Q-tip soaked in the same odorant, which caused them to habituate to that particular smell after a few presentations as determined by the substantial decrease in active sniffing. If the mice remain habituated upon presentation of a second odorant, then we conclude that they fail to differentiate the two odors. In previous studies using this paradigm, we found that odorant pairs with higher co-activation were more likely to lead to a failure to differentiate than the pairs with low co-activation. The odorants that have a complete overlap in OSN activation, therefore, should, in theory, be perceived as the same. On the other hand, the odorants that activate a large number of previously un-activated (and therefore unhabituated) OSNs should elicit a new percept.

Plotting the percentage of OSN co-activation in an odorant matrix (Fig. 6) shows the percent of OSNs activated by the odorant on the left (rows) that we’re also activated by the odorant on the top (columns). Remarkably there is a clear assymetirc relationship among some pairs that appeared to depend on the number of OSNs the odorant on the left (row) originally activated. For example, compared to CYC6-1, the most strongly and widely activating odorant in the panel, the other compounds co-activated only a subset of its activation repertoire. This is a result of the fact that in any pairwise comparison a significant number of OSNs were only activated by CYC6-1. On the contrary, from TIG4’s point of view, it was strongly co-activated by those compounds for which is it a “faithful” replacement, such as all compounds containing the tiglic moiety as their sub-structure. When we then take the odorant pair of CYC6-1 and TIG4 (Fig.6, marked in red square), there is a near complete OSN activation overlap from TIG4’s perspective, but a large number of previously un-activated OSNs from CYC6-1’s point of view. In this case, the order of odorant presentation should play a role in whether the mice can distinguish the two odors. This leads to a set of predictions about behavior based on molecular activation ranges.

**Fig. 6.**
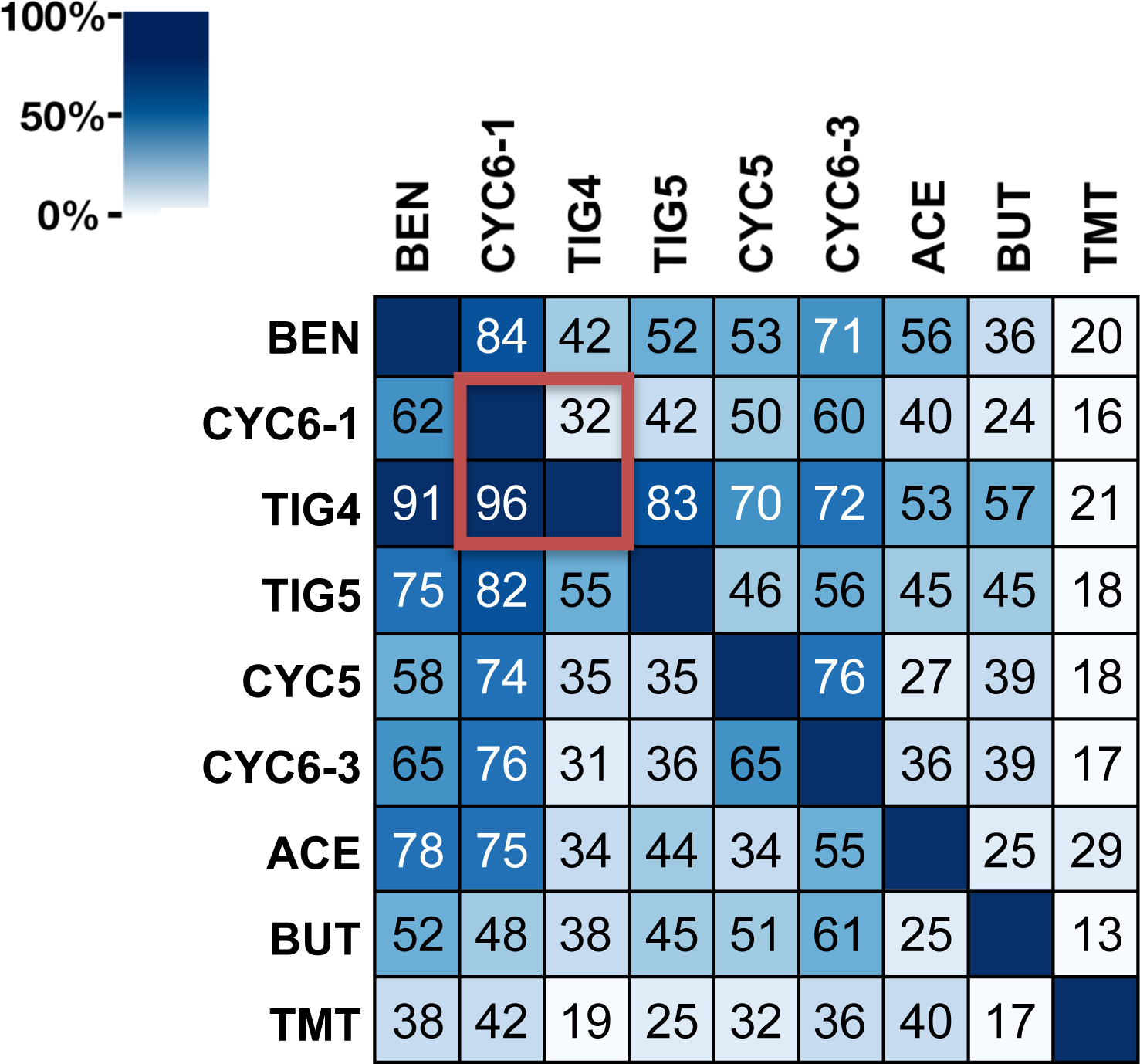
Co-activation matrix. This schematic shows the percentage of OSNs activated by the odorant on the left co-activated by the odorant on the top. A large differential between a pair of odorants indicates higher level of extraneous activation (defined in Fig. 5) in the odorant showing lower co-activation. For example, in the CYC6-1 TIG4 pair (marked by a red square), 96% of TIG4 is co-activated by CYC6-1, while the overlap is only 32% the other way around, indicative of only 4 OSNs activated by TIG4 alone and 178 OSNs activated by CYC6-3 alone. This matrix acts as a guide to understanding mouse behavior outcomes based on OSN responses (rationale explained in Fig. S5).

Based on OSN activation alone, it is possible to categorize odorant pairs in this panel into a few behavioral categories according to the co-activation matrix. Since there were no pairs that had large co-activation percentages both ways, there were no pairs in the “both way habituation” category (e.g., complete cross habituation). Pairs like CYC6-1 vs TIG4, or TIG5 vs TIG4, where there was a large differential in co-activation were placed in the likely to “habituate in one direction” category. Pairs which showed intermediate co-activation percentage were predicted to “either habituate or differentiate”. Finally, pairs that had lower than 50% co-activation both ways were predicted to “differentiate both ways”.

Indeed, odorant pairs initially put in the “habituate one way” and “differentiate both ways” categories performed as expected when tested on mice (Fig 7a, c). The prediction slightly fell apart with the “either habituate or differentiate” category odorants (Fig 7b), showing that even the small differential in % co-activation may cause asymmetric behavioral output. This has led us to consider one more variable in the prediction; OSN activation by a certain odorant in relation to all of the screened OSNs.

**Fig. 7.**
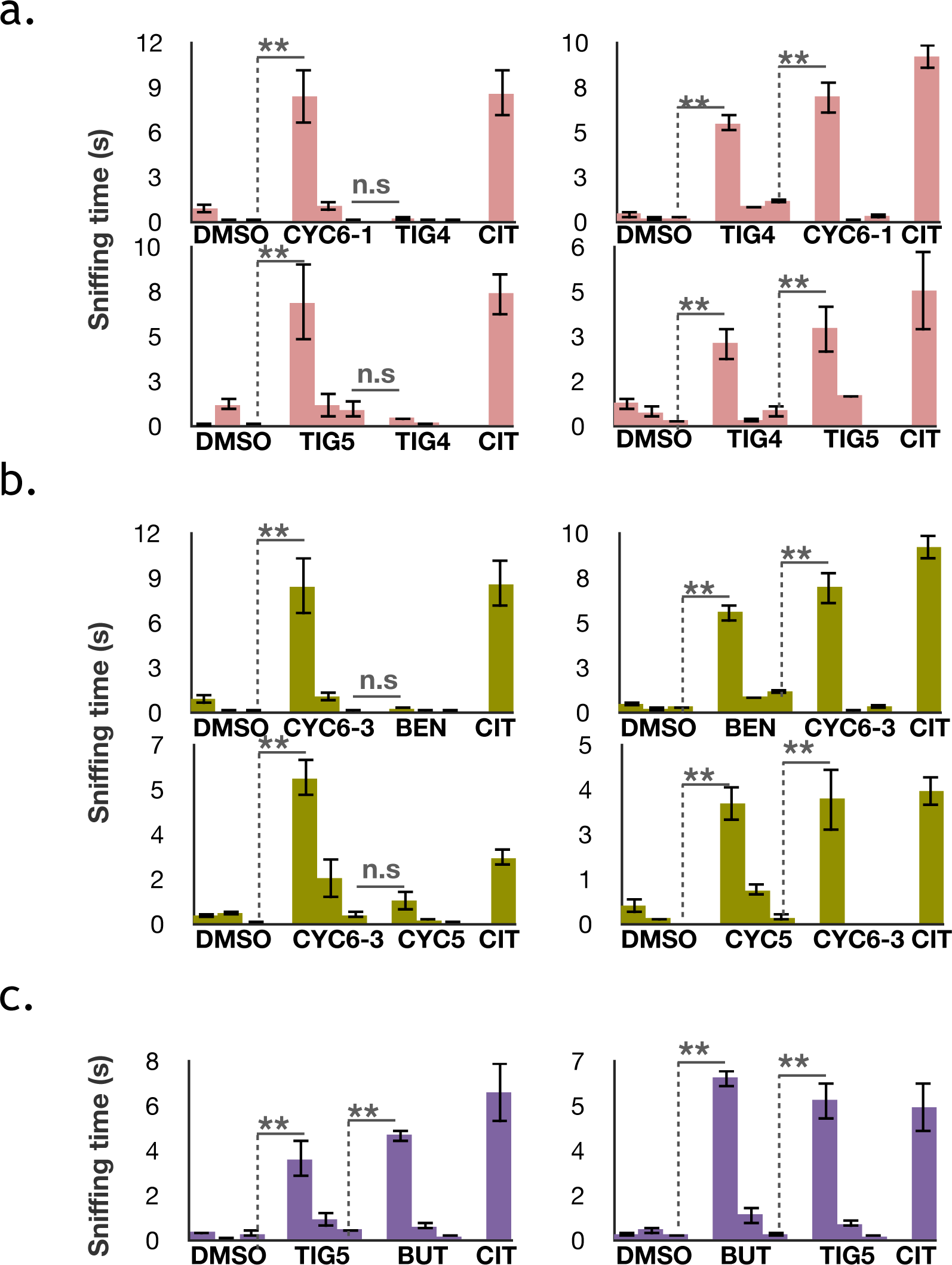
Mouse behavior results summary. Cumulative sniffing time during a 2 min odorant presentation period is represented on vertical axis, while stimulation sequence is on the horizontal. Based on the activation overlap matrix, it was possible to infer asymmetric habituation based on the sequence of odor presentation. Briefly, when there is high asymmetricity of activation overlap such as with the TIG4 and CYC6-1 pair (refer to Fig. 6), presenting the odorant with lower co-activation index first results in habituation, while the reversal in order results in differentiation. **a**. Pairs with high co-activation from the perspective of only one odorant lead to asymmetrical habituation, which corresponded to the original prediction. **b**. It was expected that pairs with with low co-activation differential would lead to either habituation or differentiation. Due to an intermediate level of activation overlap, the prediction is not as robust, and we saw both behavior types depending on the presentation order. **c**. Odorants in pairs with low overall activation overlap were differentiated both ways, as expected.

Since the predictions were initially based on the idea that full activation overlap leads to “same” percept while a large number of new OSNs activated will lead to a “different” percept from the previous habituated odor, all of the previously tested odorant pairs were plotted to reveal any correlation between OSN activation pattern and mouse behavior (Fig. 8). In this graph, the dissimilarity index (1-Jaccard index) looks at the relationship between the odorant pairs, while % new OSNs (number of new OSNs activated by the second presentation odorant divided by the total number of live OSNs screened) relates it to the global OSN activation results. When visualized this way, there seems to be a rough trend between odor pairs that lead to habituation vs those that lead to differentiation. Specifically, the presentation of the odorants with a lower Jaccard index (Fig. S2) only leads to habituation when a small number of new OSNs is activated upon the presentation of the second odorant in the pair. The slope of the differentiation border is f(x)=-.25x+.82, though pairs that sit close to the border have a smaller predictive value. Since most of the odorant pairs chosen in this study were selected to test the value of this border, we have many such points. However, most odorant pairs from the matrix do not fall close to the border; the majority would be on the top right of the graph.

**Fig. 8.**
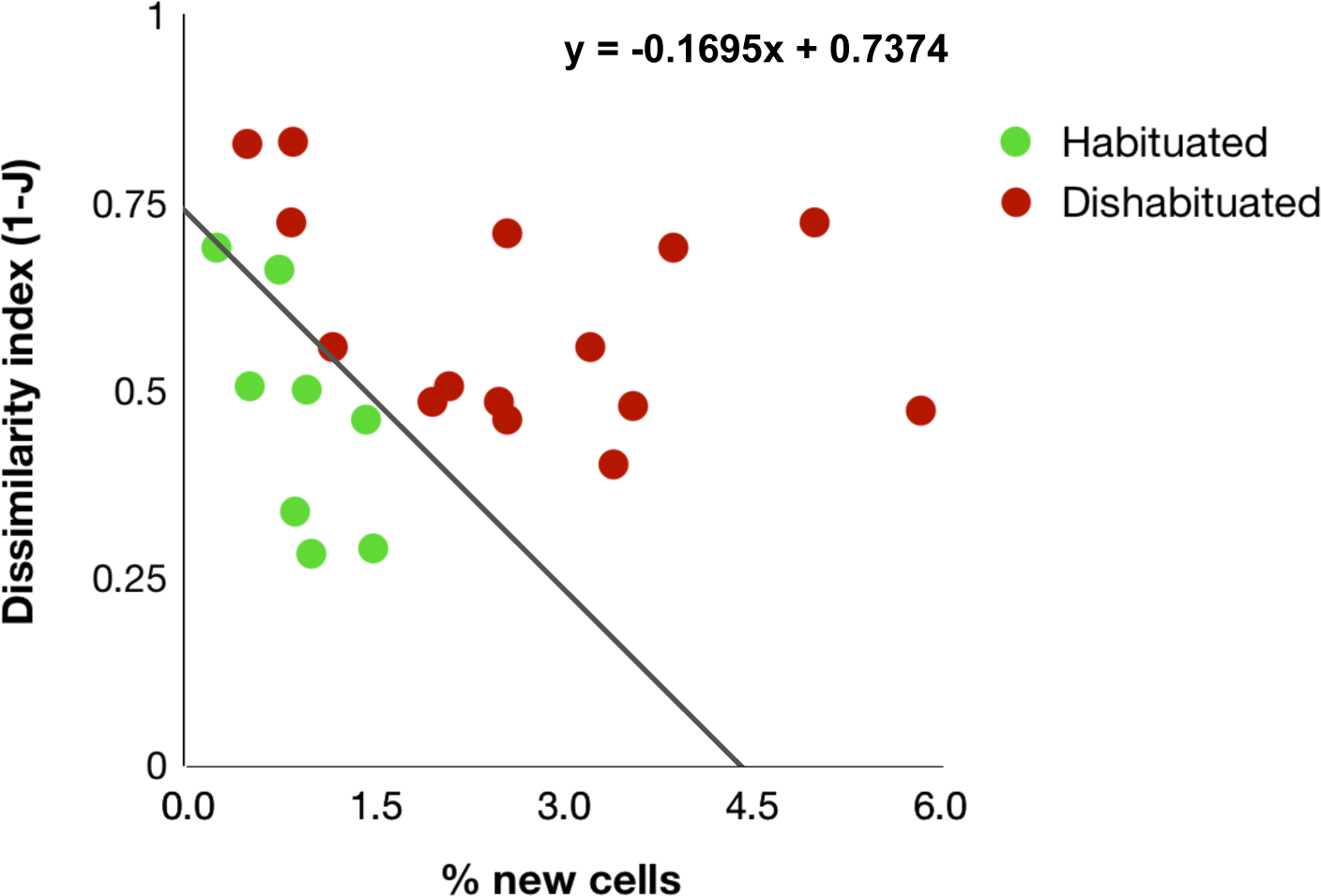
Relationship between OSN activation pattern and behavior outcomes. Summary of all behavioral experiments are plotted. The dissimilarity indices were calculated by 1-Jaccard index (from Fig. S2). % new cells is a number of OSNs that is exclusively activated by the odorant which is presented second within that pair divided by the total number of FORK+ cells screened. A decision boundary of y =-0.1695*x + 0.7374 is derived from the linear discriminant analysis using these variables. The resubstitution error is 0.0417. While the differential index is looking at pairwise comparison of odorants, the % new cells relates the odor that is presented last to global OSN activation results.

The fact that the output of the OSNs can predict odor discrimination independently of any higher level neural computation emphasizes the importance of the peripheral components of the olfactory system to forming a perception.

## DISCUSSION

Since the discovery of the Odor Receptors ORs, the olfactory field has relied on the concept of a reciprocal combinatorial code, whereby each chemical odorant can be detected by different ORs and each OR can detect a group of different chemicals [10]. Many attempts at systematic monitoring of the olfactory code have seen marginal successes for a number of reasons. First of all, there are still no solved structures of mammalian ORs to be used for high throughput computational modeling. Second, experimental validation methods such as heterologous expression still face considerable challenges. Lastly, primary chemical features of odors that allow for OR tuning have not yet been defined.

The traditional organic chemistry-based classification of odorants fails to predict biological activity, while percept-based computational analyses isolate esoteric descriptors that are difficult to chemically manipulate. Receptor level structure-activity analysis can provide a missing context to the odorant discrimination in the peripheral olfactory system. A critical finding by Manic et al [10] indicates that each mature OSN only expresses one type of OR, allowing for high throughput screening of carefully crafted odorant panels using dissociated OSN calcium imaging. Using this feature of the olfactory system, our previous work has probed how widely utilized bioisosteres in medicinal chemistry field relate to each other in terms of OR activation [3, 4, 6, 11]. Devising odorant panels using such rational method has shown great success, demonstrating OR tuning preferences among cyclic ketones based on the topological polar surface area (TPSA) descriptor [3]. Among other findings, we also saw the fluidity by which ORs may recognize esters, flipped esters, and ketones with an overall similar topological structure, which runs contrary to the widely held practice of using functional group as the main identifier of the odor grouping [4].

This study expands on the previous findings, using the concept of non-classical bio-isosterism in medicinal chemistry to show that ring odorants can be substituted by smaller non-ring fragments with retained structural integrity. Comparison of TIG4 and CYC6-1, where the activation profile of the former is almost entirely enclosed by the latter, is especially interesting, because this gives a unique insight that the retention of structural and topological profile is key to avoiding large amount of extraneous activation. By contrast, TIG5, which had an extra rotable carbon bond has an increase in non-overlapping OSN activation. BUT, which could mold into an extracted fragment of CYC6-3, showed a considerable activation in non-overlapping OSNs. These trends indicate that structural rigidity in a non-ring odorant leads to a more faithful replacement. This may have significant implications for drug design, where ring structure may need to be removed for pro-drug generation while avoiding activation of non-target receptors.

Our findings with ACE and BEN activation overlap recapitulate the previous observation that functional group identity may not be a key distinguishing feature in odorants with the similar topological profile [4]. Such significant overlap between aldehydes and ketones, as well as ketones and esters, and on large scale screens, suggests strong link between odorants with different polar functional groups, and urges for further exploration of such link. Since our findings with heteroaromatic ketones suggest a TPSA-based trend in odorant-OR interaction [3], perhaps this variable could be a starting point for a large-scale probing of the relationship among the functional groups in structurally analogous compounds.

## MATERIALS AND METHODS

### Chemicals

The panel odorants were designed around the lead aldehyde odorant, benzaldehyde (BEN), consisting of cyclohexene-1-carboxaldehyde (CYC6-1), tiglinaldehyde (TIG-4), 2-methyl 2-pentenal (TIG-5), cyclopentane carbaldehyde (CYC-5), 3-cyclohexene-1-carbaldehyde (CYC6-3), and butyraldehyde (BUT). Acetophenone (ACE), a ketone was added to understand the potential link between aldehyde and ketone activation spaces. 2,3,5-trimethyl-3-thiazoline (TMT) was added to the panel as an outgroup and purchased from Clontech Enterprices. The rest of the odorants were sourced from Sigma-Aldrich. On the day of the imaging experiments, the odorant stocks were reconstituted in >99% DMSO (Sigma-Aldrich), and subsequently diluted 1000-fold in Ringer’s solution to make up a 30uM stimulation solution.

### Animal and tissue collection

All animal procedures conformed to the guidelines for care and use of animals of Columbia University and were reviewed and approved by the Institutional Animal Care and Use Committee. OMP-Cre–driven GCaMP6f mice used in this work were generated by crossing the OMP-Cre line (#006668, the Jackson Laboratory) with the line B6;129S-Gt(ROSA)26Sortm95.1(CAG-GCaMP6f)Hze/J (#024105, the Jackson Laboratory). In these compound mutant mice, the expression of the genetically encoded calcium sensor GCaMP6f was restricted to the mature OSNs. All mice were reared and maintained in the department animal facility. OSNs were isolated from 5- to 8-week-old OMP-Cre–driven GCaMP6f male mice with a genotype of OMP-Cre+/−GCaMP6f−/−. The mice were overdosed with anesthetics [ketamine (90 mg kg−1, intraperitoneally) and xylazine (10 mg kg−1, intraperitoneally)] and decapitated. The head was cut open sagittally, and the septum was removed to expose the medial surface of the olfactory epithelium and turbinates. The olfactory epithelium and turbinates were dissected and collected in divalent-free Ringer’s solution [145 mM NaCl, 5.6 mM KCl, 10 mM Hepes, 10 mM glucose, and 4 mM EGTA (pH 7.4)]. The tissue was incubated at 37°C for 45 min in 5 ml of divalent-free Ringer’s solution containing collagenase (0.5 mg ml−1), bovine serum albumin (5 mg ml−1; Sigma-Aldrich), dispase (5 U ml−1; Roche), and deoxyribonuclease II (50 mg ml−1; Sigma-Aldrich). The tissue was then transferred to a clean tube of culture medium and washed using the previously warmed culture medium. Culture medium consisted of DMEM (Dulbecco’s modified Eagle’s medium)/F12 (Gibco BRL) supplemented with 10% fetal bovine serum, 1× insulin-transferrin-selenium (Gibco BRL), penicillin (100 U ml−1) and streptomycin (100 mg ml−1; Gibco BRL), and 100 mM ascorbic acid (Sigma-Aldrich). The OSNs were dissociated by tapping the tube containing the tissue. The OSNs (50-µl volume) were split onto four concanavalin-coated glass coverslips (10 mg ml−1; Sigma-Aldrich) and placed in 35-mm petri dishes. After allowing the cells to settle for 20 min, 2 ml of culture medium was added to each dish, and the dishes were placed at 37°C for at least 1 hour.

### Calcium-imaging of the dissociated OSNs

After being washed with fresh Ringer’s solution, the coverslips were mounted on a recording chamber. Imaging was carried out at room temperature on an inverted fluorescence microscope (IMT-Olympus) equipped with an SIT camera (C10600, Hamamatsu Photonics), a Lambda XL light source (Sutter Instrument), and Lamba-10B optical filter changer (Sutter Instrument). Using a 1260 Infinity HPLC system (Agilent Technologies), the dissociated OSNs were stimulated with the odorants in random order. A final stimulation with a 10 µM forskolin (Sigma-Aldrich) solution was made to assess the viability of the OSNs. Recordings were made at 490-nm excitation and 520-nm emission. Images were taken every 4 s, and there was a 4-min delay between odorant stimulations. The cell fluoresncence from the images was then quantified using MetaMorph Premier software (Molecular Devices LLC). Cells exhibiting an intensity increase of at least 10% Δ*F*/*F*0 amplitude increase over baseline between 8 and 12 frames after the odorant injection were considered responsive cells.

### Data analysis of calcium imaging recording

Data were analyzed using custom Matlab(MathWorks) scripts. For molecular descriptors-based clustering, A total of 1666 molecular descriptors for the panel odorants were downloaded through e-Dragon free applet (www.vcclab.org). All descriptors were *z*-scored before used for clustering. Dendrograms were then generated based on Euclidean distances between odors. For clustering of biological results, neuronal responses were scored with binary values, where “1” indicates “response,” and “0” indicates “no response.” The Pearson correlation coefficients between odors were generated with Matlab build-in function *corrcoef*, then clustered based on their Euclidean distances. To evaluate the potency of each molecular descriptor in predicting neuronal responses to different odors, a Spearman’s rank correlation index between each single descriptor-based clustering result and OSN response clustering result. The indices were then ranked to identify descriptors that best recapitulate features of OSN responses.

### Habituation-dishabituation behavioral test

Similarities in perceptual odor quality among the panel of odorants were evaluated by a habituation-dishabituation olfactory test in the mouse. Thirty minutes before experimentation, 5- to 8-week-old OMP-Cre+/− GCaMP6f−/− male mice were placed individually into a hood in an empty mouse cage containing a cotton swab soaked in DMSO/Ringer’s solution (1:1000). Each animal was then stimulated three consecutive times over 2 min with the DMSO/Ringer’s solution soaked in a cotton swab as a negative control. Then, they received three consecutive presentations of a cotton swab soaked in the first odorant solution at 30 µM. Each presentation lasted 2 min, with a 1-min interval between presentations. Following a 1-min rest, animals were then given three presentations of the second odor in a similar manner. Following a final 1-min break, a 30 µM solution of citral was given in a 2-min single stimulation as a positive control. The cumulative sniffing time of the cotton swab was recorded using a silent clock. An ANOVA statistic comparison, followed by post hoc paired *t* test, was performed on the results using StatView. Each mouse was used only once with the same odorant. Mice that were unable to detect the first odorant stimulation or that responded to the negative control were removed from further analysis.

### Linear discriminant analysis of behavior results vs OSN responses

The relationship between the OSN activation and mouse behavior output was plotted using dissimilarity index (1-Jaccard index) and % new cells (#neurons recruited exclusively by the second odorant divided by number of FORK+ cells). The dissimilarity indices and cell recruitment percentage numbers were then used as predictors for linear discriminant analysis using matlab build-in function *fitcdiscr*. Data points were either classified as habituation (0) or differentiation (1). After Bayesian optimization, a model was generated to predict future data. A decision boundary was plotted according to this model.

---

**Table S1.** A list of top eDRAGON descriptors. This table lists the molecular descriptors acquired from eDRAGON applet which best recapitulate the hierarchical clustering of the odorants based on OSN responses. *Rho* is a Spearman rank coefficient, *name* is descriptor identifier within the eDRAGON repertoire.

| **Rho** | **Name** | **Full name** | **Descriptor type** |
| --- | --- | --- | --- |
| 0.802 | H-051 | H attached to alpha-C | Atom-centred fragment |
| 0.781 | ASP | Asphericity | Geometrical descriptor |
| 0.779 | EEig01x | Eigenvalue 01 from edge adj. matrix weighted by edge degrees | Eigenvalue |
| 0.779 | ESpm15x | Spectral moment 15 from edge adj. matrix weighted by edge degrees | Edge adjacency index |
| 0.773 | ESpm14x | Spectral moment 14 from edge adj. matrix weighted by edge degrees | Edge adjacency index |
| 0.766 | Ram | Ramification index | Topological index |
| 0.766 | ESpm03u | Spectral moment 03 from edge adj. matrix | Edge adjacency index |
| 0.748 | ESpm11x | Spectral moment 11 from edge adj. matrix weighted by edge degrees | Edge adjacency index |
| 0.748 | ESpm12x | Spectral moment 12 from edge adj. matrix weighted by edge degrees | Edge adjacency index |
| 0.747 | ESpm13x | Spectral moment 13 from edge adj. matrix weighted by edge degrees | Edge adjacency index |

**Fig. S2.**
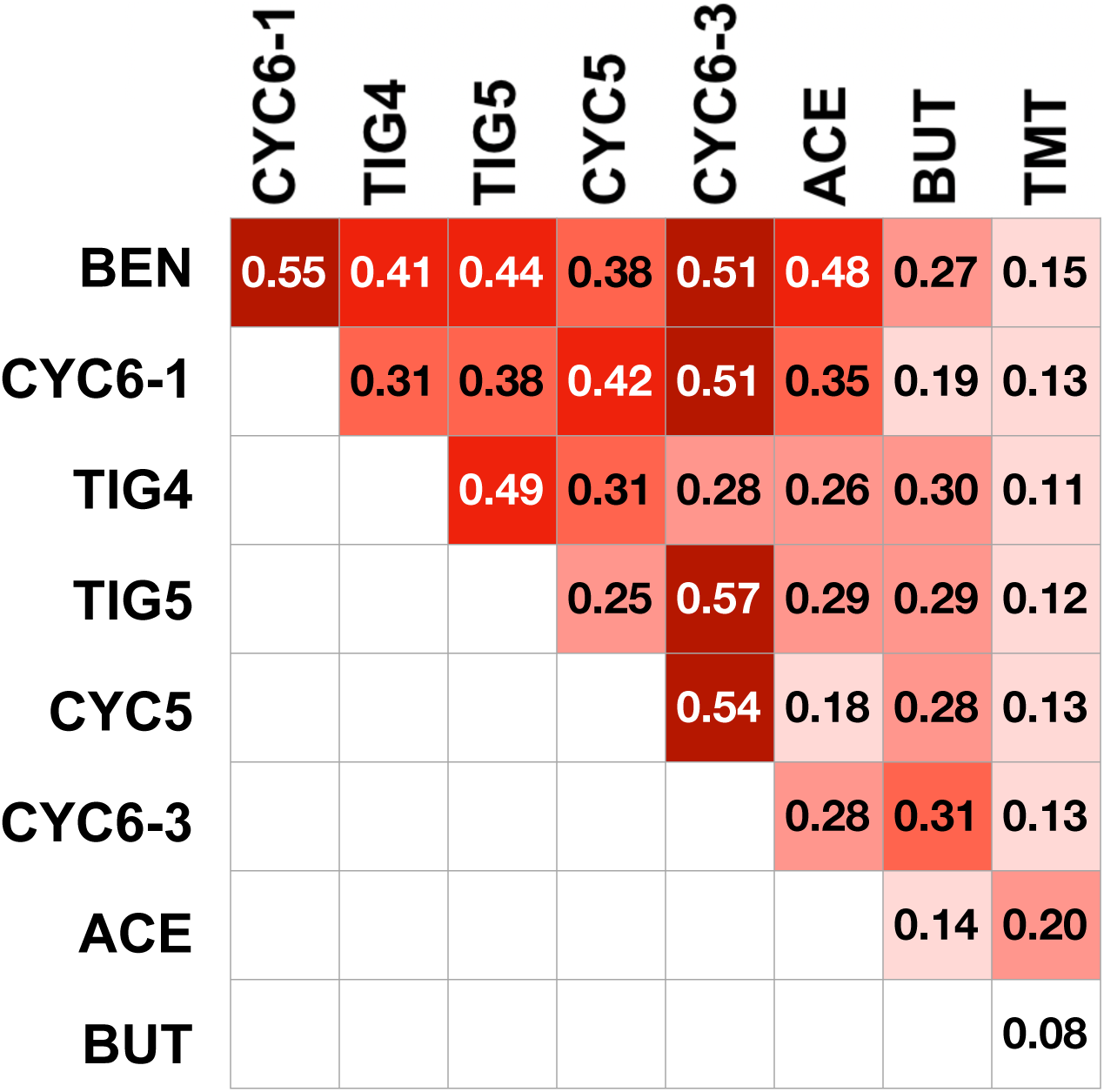
Jaccard similarity index table for the odorant pairs. Jaccard index is used to measure the similarity between the responding OSN population in odorant pairs, and is defined as the size of the intersection divided by the size of the union of the sample sets. This index is used in Fig. 8 for pairwise comparison of the odorants.

**Fig. S3.**
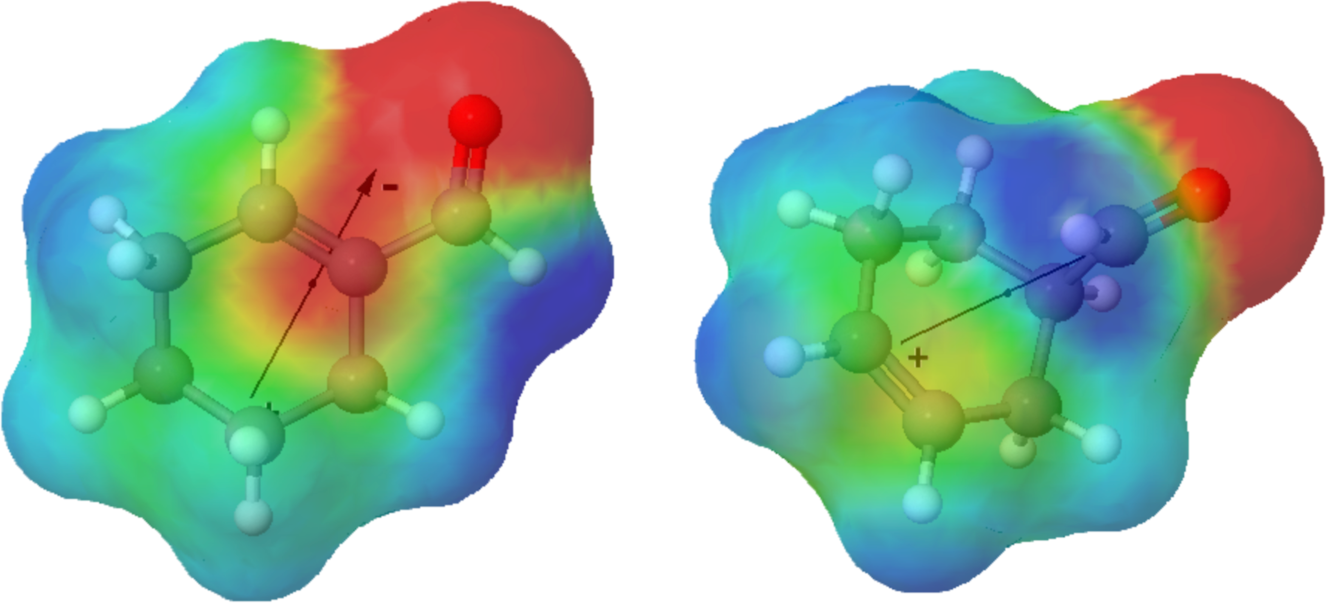
3D comparison of the 6-membered alkyl rings of the panel. While both CYC6-1 and CYC6-3 possess the same chemical formula and the TPSA, the electronegativity distribution and the difference in afforded structural freedom leads to a large number of non-overlapping cells among the two.

**Fig. S4.**
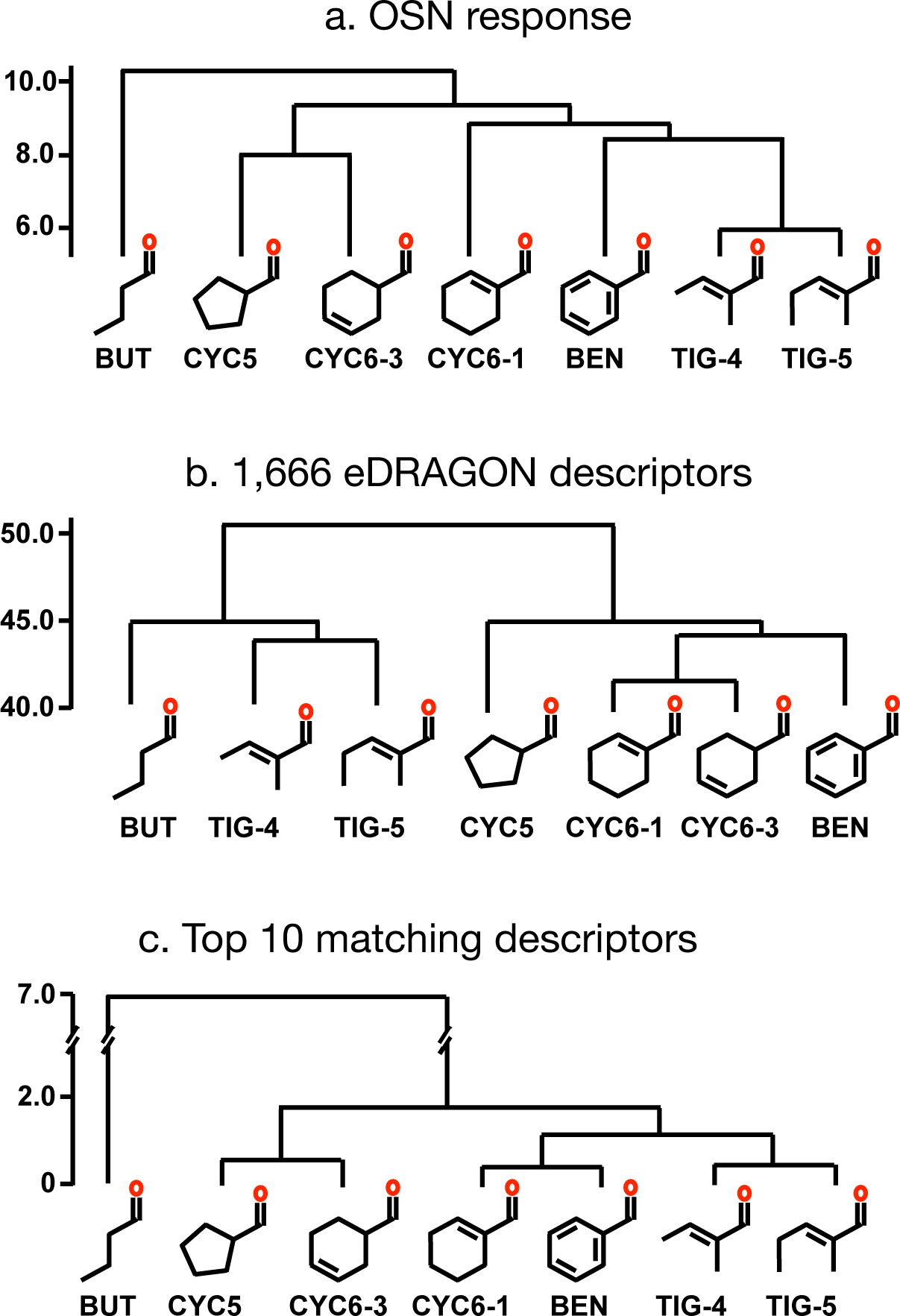
Hierarchical cluster analysis of the panel’s aldehydes. Aldehydes clustered according to: **a.** Biological response similarity based on calcium imaging of dissociated OSNs, **b.** Chemical similarity as evaluated by all 1666 molecular descriptors downloaded through the e-Dragon applet, **c**. Top 10 ranking eDRAGON descriptors listed in Table S1.

**Fig. S5.**
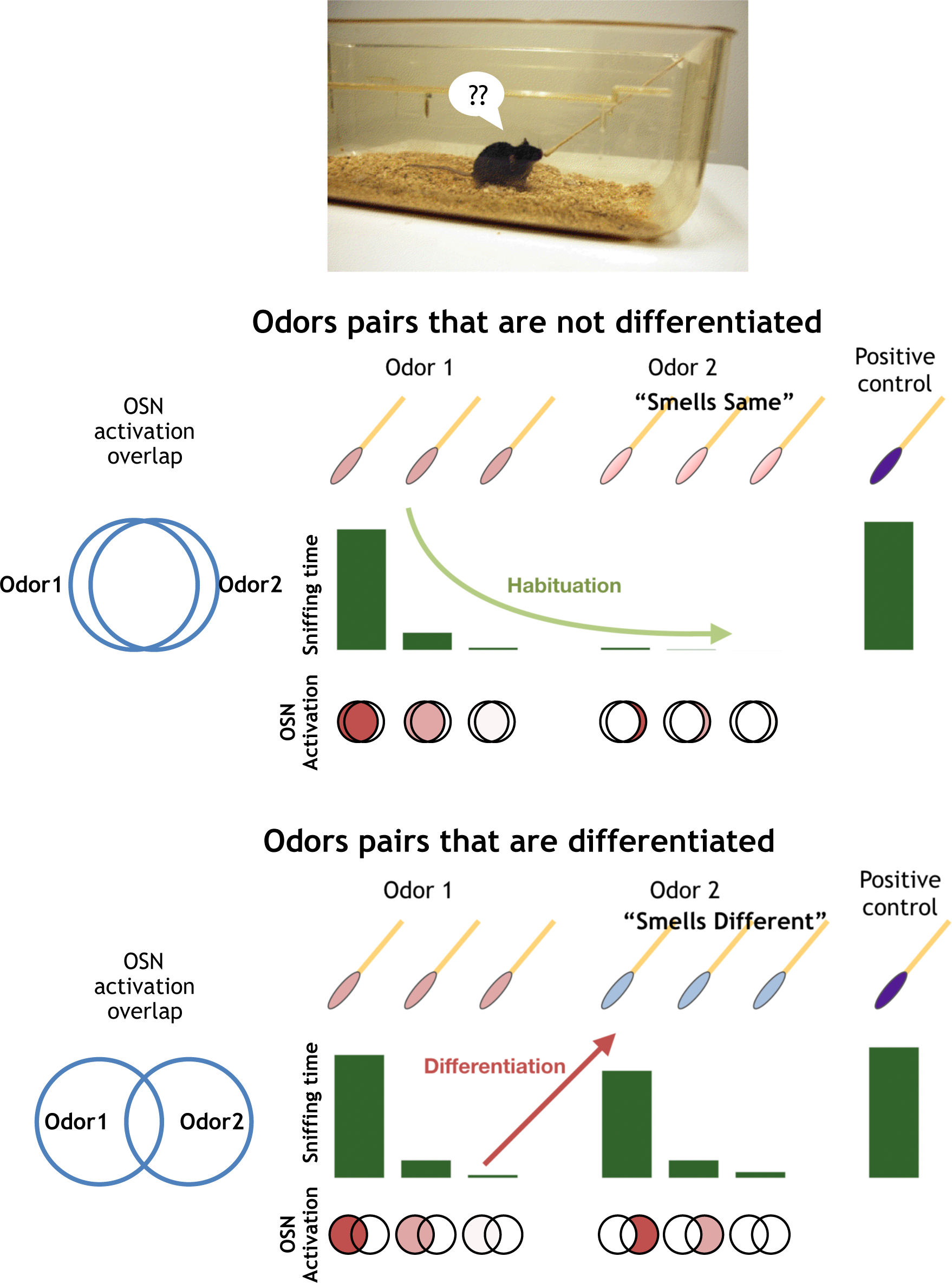
Rationale for mouse behavior experiments, as well as potential link between OSN activation pattern and the behavioral output. **a.** The odor pairs with high level of overlap, where odor which is presented second does not exclusively recruit a large number of OSNs. It is expected that mice would fail to differentiate such odors. The number of OSNs recruited exclusively is plotted as “% new cells” against Jaccard index in Fig. 8. High level of overlap, such as one exemplified on the left hand schematic, combined with low “% new cells” is more likely to lead to continued habituation of the mouse to the second odorant. **b.** Conversely, the mice are more likely to differentiate the odor pair with a low Jaccard index or a large “% new cells” at odor #2 presentation.

